# Efficient ageing: Simulated lesion of the structural connectome reveals optimised decline in the healthy ageing brain

**DOI:** 10.64898/2026.05.29.728718

**Authors:** N. Kudriavtsev, M. Rosso, G. Fernández-Rubio, E. Serra, M. L. Kringelbach, P. Vuust, L. Bonetti

## Abstract

Healthy ageing is associated with widespread white-matter change and altered connectome organisation, yet the link between local microstructural decline and whole-brain network communication remains unclear. Here we combined tract-based spatial statistics (TBSS) with probabilistic tractography and graph analysis to quantify the connectome-level consequences of age-sensitive white-matter hotspots. In two independent diffusion MRI datasets of healthy young and older adults (n = 144 total; Dataset 1: 77 participants; Dataset 2: 67 participants), we first identified age-sensitive fractional anisotropy clusters and then used them as constraints in a cross-dataset simulated-lesion framework. This allowed us to estimate how strongly each structural connection depended on age-sensitive tissue and to test the resulting network effects against matched-mass null lesions. Across both datasets, ageing hotspot lesioning reduced global efficiency, but consistently less than expected under the null model, indicating a less-than-random disruption of network integration. Age-sensitive tissue was disproportionately embedded in the brain’s integrative backbone: proportional degree loss was strongest in high-degree nodes, rich-club connections showed the greatest hotspot dependence, and nodewise losses were concentrated in frontal, cingulate and subcortical association systems, whereas posterior sensory and temporo-limbic regions were relatively spared. These findings suggest that healthy ageing reflects a selective and constrained reconfiguration of structural connectivity rather than simply pointing to uniform decline.

## Introduction

Population ageing is accelerating worldwide, and with it the prevalence of late-life cognitive change and neurodegenerative disease (United Nations DESA, 2023; GBD 2019 Dementia Forecasting Collaborators, 2022; Prince et al., 2024). Understanding how ageing reshapes brain organisation is therefore both a scientific and public-health priority (Beard et al., 2016; Sala-Llonch et al., 2015; Sabayan et al., 2023). A central message from contemporary cognitive neuroscience is that healthy ageing is not a single process, but a constellation of biological and physiological changes that unfold across levels of description – from cellular and microstructural alterations to large-scale reconfiguration of brain systems (Gunning-Dixon et al., 2009; Sexton et al., 2014; Jones et al., 2013; Deery et al., 2023). In parallel, influential frameworks emphasise the tension between vulnerability and resilience: some older adults maintain brain integrity, others tolerate neural change with less behavioural impact, and still others show compensatory reorganisation (Cabeza et al., 2018). Together, these perspectives motivate integrative approaches that can connect microstructural ageing to the network-level constraints that ultimately shape cognition (Aerts et al., 2016; Riedel et al., 2022), which is a core goal of this study.

One of the most consistent structural signatures of healthy ageing is change in cerebral white matter (WM). Diffusion MRI (dMRI) studies report widespread age-related reductions in fractional anisotropy (FA) – a measure of the directionality of water diffusion in tissue, often interpreted as marker of fibre coherence and/or myelin integrity – with prominent effects in association pathways (Gunning-Dixon et al., 2009; Sexton et al., 2014; Barrick et al., 2010; Poulakis et al., 2021). These WM differences are not merely descriptive: they relate to behavioural variability, help explain why cognitive ageing trajectories vary across individuals (Gunning-Dixon & Raz, 2000; Coelho et al., 2021; Deery et al., 2023), and may be sensitive to age-related pathophysiology and cognitive status, including in preclinical and prodromal Alzheimer’s disease (Sexton et al., 2014; Peter et al., 2025). A widely used approach to localising these WM differences and mapping robust age effects across the lifespan is tract-based spatial statistics (TBSS), which projects dMRI metrics onto a common WM skeleton and enables group comparisons (Smith et al., 2006; Sexton et al., 2014; Poulakis et al., 2021). However, TBSS alone does not tell us which structural connections are most constrained by a given age-sensitive cluster, nor how that constraint affects large-scale properties of the brain.

Another piece of the puzzle – systems-level perspective – frames ageing as a gradual reconfiguration of the structural connectome. Healthy brain networks typically exhibit a small-world topology that balances integration and segregation (Latora & Marchiori, 2001; Rubinov & Sporns, 2010). Within this architecture, highly connected hubs form a rich-club – a metabolically costly backbone that supports long-range communication across systems and large-scale neural populations (van den Heuvel & Sporns, 2011; Rubinov & Sporns, 2010). The literature indicates that this backbone is disproportionately vulnerable: targeted damage to hub-rich routes reduces global efficiency more than a random lesion matched in size, motivating network-based approaches to study brain vulnerability (Aerts et al., 2016; van den Heuvel & Sporns, 2011). In ageing, connectome studies often report reduced global efficiency, altered clustering, and changes in hub-related organisation – patterns that broadly mirror the vulnerability of long-range association pathways highlighted by diffusion studies (Zhao et al., 2015; Riedel et al., 2022; Stumme et al., 2022). Large-sample work further suggests that ageing does not affect all network elements uniformly, but shows system-specific footprints and prominent effects in integrative routes that are especially relevant for cognitive function (Madole et al., 2021; Li et al., 2023). However, despite these coherent findings, microstructural and large-scale connectome changes in healthy ageing are usually studied independently. In other words, the field has detailed maps of where WM ageing occurs, and a growing literature on how network topology changes with age, but the bridge between these levels is still missing.

In the present work, we attempt to connect microstructural ageing to network communication in a mechanistic and interpretable way. We first identify age-sensitive WM hotspots using TBSS, and then use these hotspots as constraints during subject-level probabilistic tractography to quantify how strongly each structural connection depends on age-sensitive tissue (Smith et al., 2006; Tzourio-Mazoyer et al., 2002; Behrens et al., 2003, 2007). This yields an individualised, cluster-proportional simulated lesion, enabling direct tests of how hotspot-constrained weakening changes network integration and local organisation (Rubinov & Sporns, 2010). To ensure biologically plausible tractography-derived connectomes, we apply consistency-based thresholding (Roberts et al., 2017), and to interpret effect sizes conservatively we benchmark observed network impacts against matched-mass null models. Across two independent datasets, this framework allows us to ask not simply whether ageing-related white-matter change reduces structural connectivity, but whether the observed pattern of decline is especially disruptive – or instead constrained in a way that preserves network-level communication capacity. In this sense, the study aims to bridge microstructural maps, connectome topology, and influential theories of ageing centred on disconnection, compensation, and resilience.

## Results

We combined TBSS-defined white-matter hotspots with subject-level probabilistic tractography to estimate how strongly each structural connection depended on age-sensitive tissue, and then used a simulated-lesion framework to test the network consequences of that dependence (Figure 1). The final sample comprised 144 healthy participants across the two datasets, including 37 young (mean age = 21.89 years, s.d. = 2.05) and 40 older adults (mean age = 67.50 years, s.d. = 5.46) in Dataset 1, and 41 young (mean age = 22.37 years, s.d. = 2.57) and 26 older adults (mean age = 68.77 years, s.d. = 5.30) in Dataset 2. The two-dataset cross-design allowed hotspot definition and hotspot-constrained tractography to be separated across samples, avoiding circularity. We first examined effects on global efficiency, then tested whether highly connected nodes were disproportionately affected, followed by nodewise analyses of degree and clustering coefficient, and finally evaluated whether the ageing hotspots were preferentially embedded in rich-club routes, all of which provide a comprehensive evaluation of network integration.

**Figure 1:**
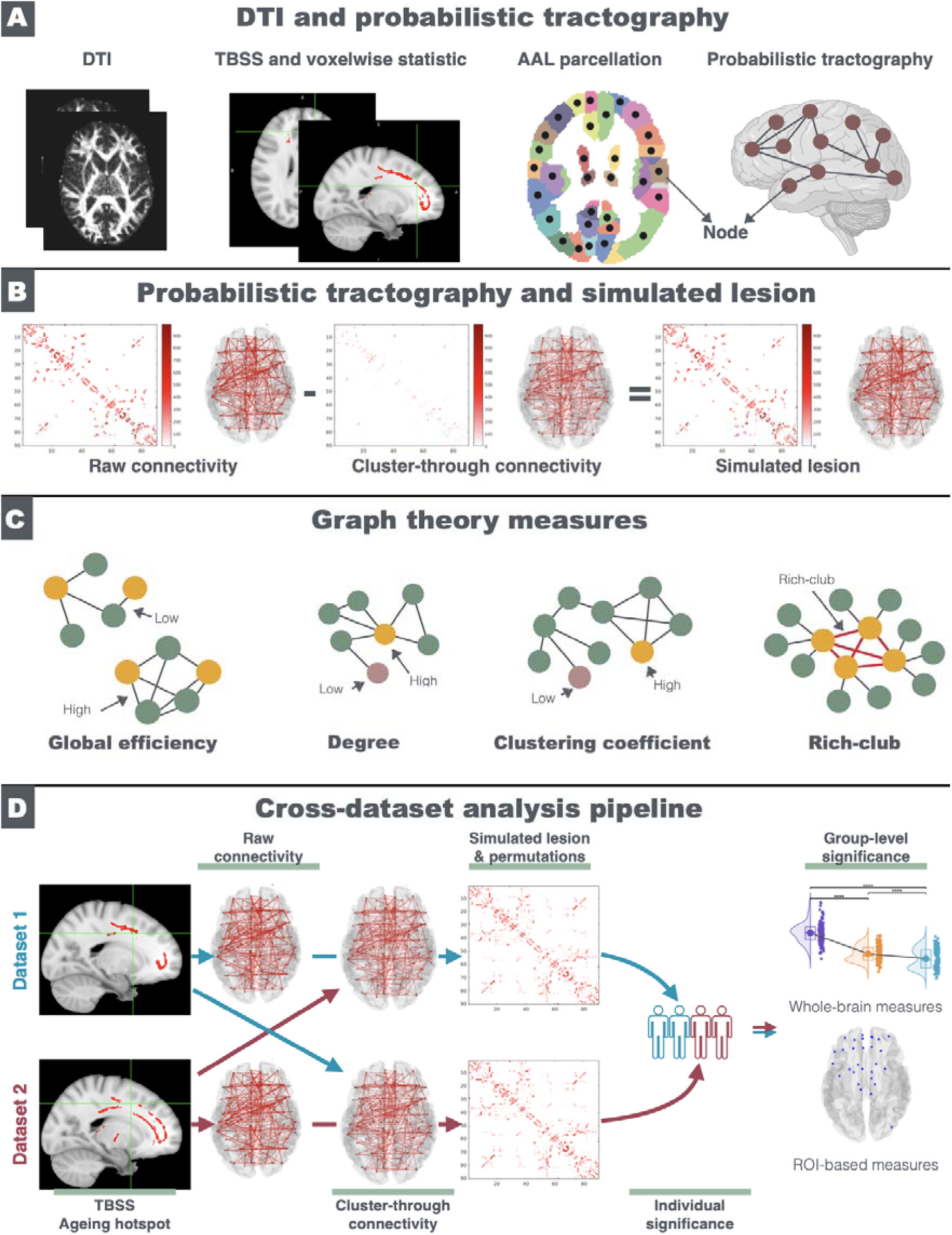
Cross-dataset simulated-lesion framework linking age-sensitive white matter to connectome-level consequences. **(A)** Diffusion MRI preprocessing and hotspot definition. For each dataset, diffusion data were preprocessed, fractional anisotropy (FA) maps were analysed with TBSS, and significant older-versus-younger white-matter clusters were identified with voxelwise statistics and cluster-based permutations. These age-sensitive clusters were then related to an AAL-90 parcellation and probabilistic tractography. **(B)** Implementation of the simulated lesion. For each participant, a raw structural connectivity matrix was first reconstructed without any waypoint constraint. A second “cluster-through” matrix was then reconstructed by counting only streamlines that passed through the age-sensitive hotspot. Subtracting the cluster-through contribution from the raw matrix yielded an individual, cluster-proportional simulated lesion. **(C)** Graph-theoretical readouts. The network consequences of the simulated lesion were quantified with whole-brain and nodewise graph measures, including global efficiency, degree, clustering coefficient, and rich-club organisation. **(D)** Cross-dataset analysis pipeline. Age-sensitive hotspots were identified separately in each dataset, but used as tractography waypoints in the other dataset to reduce circularity. Whole-brain effects were tested at the participant level against matched-mass null models, whereas nodewise effects were combined across datasets to identify regions showing consistent above-or below-chance loss at the group level.

### Global efficiency

Hotspot-constrained lesioning produced a clear but less-than-expected reduction in global efficiency. Based on the participant-level permutation results, the empirical efficiency drop was smaller than the matched-mass null expectation in all participants across the combined sample (166/166 participants; binomial p = 1.07e-50), with the same pattern present in both Dataset 1 (78/78; p = 3.31e-24) and Dataset 2 (88/88; p = 3.23e-27). Thus, although ageing-sensitive white matter reduced network integration, the observed loss was consistently less disruptive than a size-matched random lesion (**Figure 2**).

**Figure 2:**
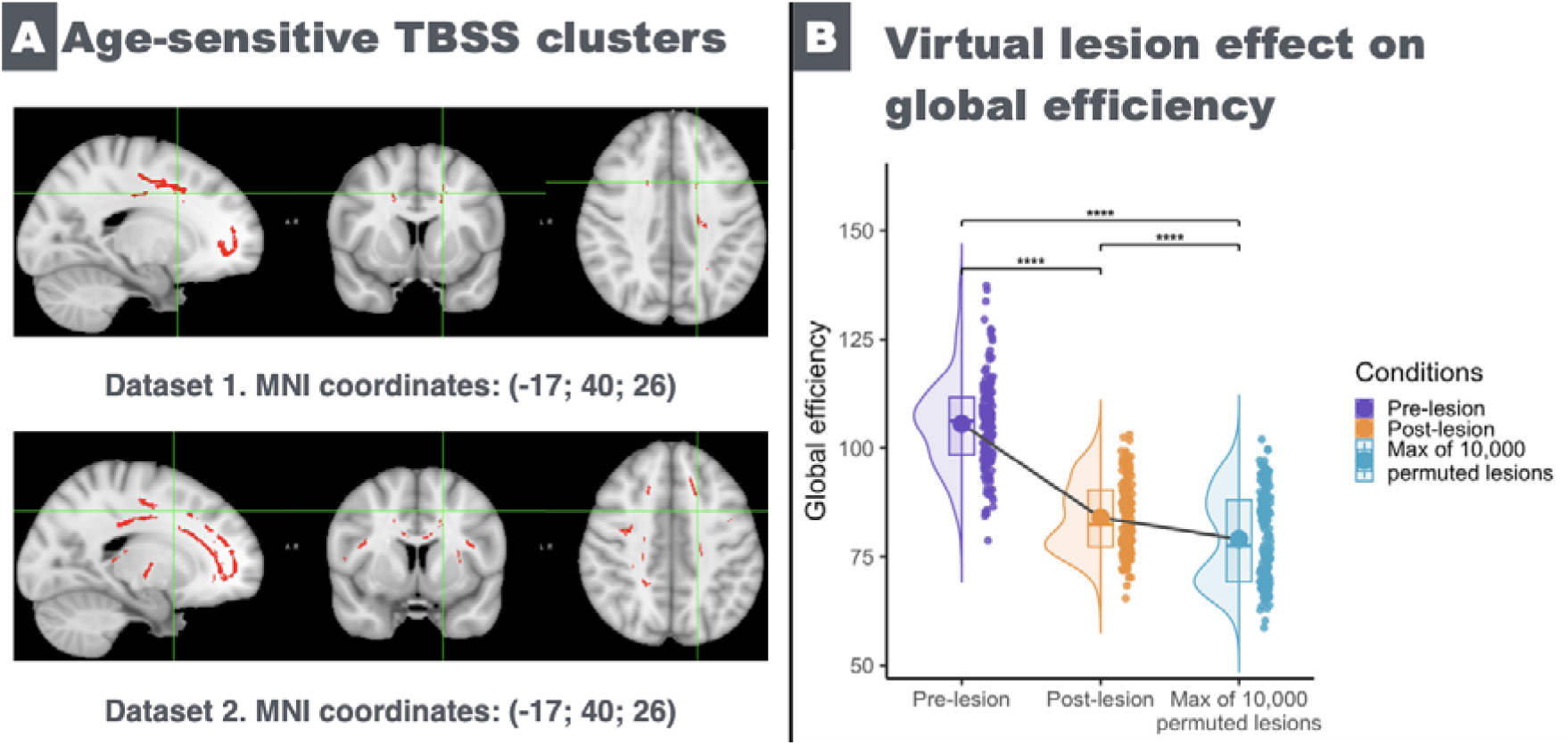
Age-sensitive TBSS clusters and their effect on global efficiency. **(A)** Significant TBSS clusters showing lower FA in older than younger adults in Dataset 1 and Dataset 2, displayed on representative sagittal, coronal, and axial slices in MNI space. These clusters served as the age-sensitive white-matter hotspots for the simulated-lesion analysis. **(B)** Effect of the simulated lesion on global efficiency at the main 30% consistency threshold. Violin/box plots show pre-lesion global efficiency, postlesion global efficiency, and the upper end of the matched-mass null distribution derived from 10,000 permuted lesions. Across both datasets, hotspot-constrained lesioning reduced global efficiency, but the observed reduction was smaller than expected under the size-matched random-lesion model, consistent with a less-than-expected disruption of network integration. Dots indicate individual participants; boxplots show the median and interquartile range.

### Hub-disruption analysis

This non-random pattern was further reflected in the hub-disruption analysis. The hub-disruption index (HDI) was significantly positive across the full sample (mean slope = 0.0174, s.d. = 0.0162, 95% CI [0.0149, 0.0199], t(165) = 13.82, p = 1.15e-29), indicating that nodes with higher baseline degree showed stronger proportional loss after lesioning (**Figure 3A**). The effect was present in both datasets separately (Dataset 1: mean = 0.0271, s.d. = 0.0152, t(77) = 15.78, p = 3.90e-26; Dataset 2: mean = 0.00888, s.d. = 0.0118, t(87) = 7.05, p = 2.03e-10).

**Figure 3:**
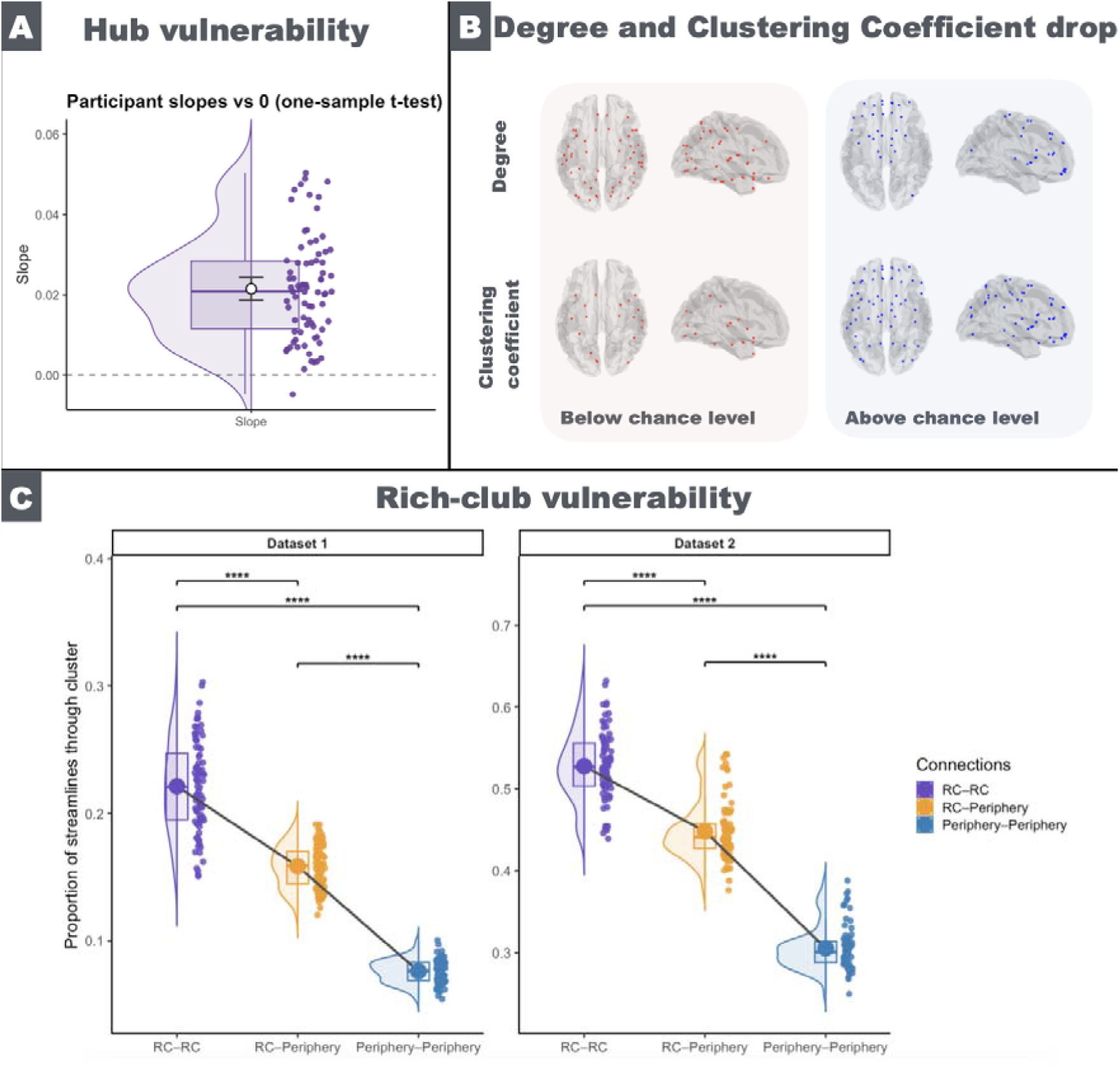
Hub, nodewise, and rich-club vulnerability of the structural connectome. **(A)** Hub vulnerability at the main 30% consistency threshold. Participant-wise hub-disruption slopes are shown relative to zero; positive values indicate that nodes with higher baseline degree showed stronger proportional loss after simulated lesioning. The dashed line marks zero slope. **(B)** Nodewise effects on degree and clustering coefficient. Regions showing below-chance loss (left) and above-chance loss (right), relative to the matched-mass permutation null, are displayed for weighted degree and clustering coefficient. Group-level maps were obtained by pooling participant-level permutation results across datasets and testing consistency of effect direction across individuals. **(C)** Rich-club vulnerability in Dataset 1 and Dataset 2. Violin/box plots show the proportion of streamline weight routed through the age-sensitive hotspot for rich–rich, rich–periphery, and periphery–periphery connections at 30% density. In both datasets, hotspot dependence followed a graded pattern, with the strongest dependence in rich–rich connections, intermediate dependence in rich–periphery connections, and the lowest dependence in periphery–periphery connections. Dots indicate individual participants; boxplots show the median and interquartile range.

### Nodewise degree and clustering coefficient

At the regional level, degree loss was not spatially uniform (**Figure 3B**). Above-chance degree loss was concentrated in frontal, cingulate, and subcortical integrative territories (29 regions), including bilateral superior and orbitofrontal cortex, medial frontal/vmPFC regions, supplementary motor/paracentral areas, anterior and mid-cingulate cortex, and bilateral striatal-thalamic regions. Specifically, the regions showing above-chance degree loss were: *L Precentral, L Frontal Sup, L Front Sup Orb, L Front Mid, L Front Mid Orb, L Front Inf Ope, L Front Inf Tri, L Supp Motor Ar, L Front Sup Med, L Ventr Med PFC, L Cingulum Ant, L Paracentr Lob, L Caudate, L Putamen, L Pallidum, L Thalamus, R Thalamus, R Putamen, R Caudate, R Paracentr Lob, R Occipital Mid, R Cingulum Mid, R Cingulum Ant, R Ventr Med PFC, R Front Sup Med, R Supp Motor Ar, R Front Mid Orb, R Front Sup Orb, and R Frontal Sup*.

By contrast, below-chance degree loss was concentrated in posterior sensory and temporo-limbic/peripheral systems (49 regions), including bilateral calcarine, cuneus, lingual, occipital, fusiform, postcentral, superior/inferior parietal, supramarginal/angular, Heschl, superior/middle/inferior temporal and temporal-pole regions, together with bilateral parahippocampal and amygdalar regions. The full set of regions showing below-chance degree loss was: *L Rolandic Oper, L Subgenual, L Front Med Orb, L Insula, L Cingulum Post, L ParaHippocamp, L Amygdala, L Calcarine, L Cuneus, L Lingual, L Occipital Sup, L Occipital Mid, L Occipital Inf, L Fusiform, L Postcentral, L Parietal Sup, L Parietal Inf, L SupraMarginal, L Precuneus, L Heschl, L Temporal Sup, L Tempr Pol Sup, L Tempr Pol Mid, L Temporal Inf, R Temporal Inf, R Tempr Pol Mid, R Temporal Mid, R Tempr Pol Sup, R Temporal Sup, R Heschl, R Angular, R SupraMarginal, R Parietal Inf, R Parietal Sup, R Postcentral, R Fusiform, R Occipital Inf, R Lingual, R Cuneus, R Calcarine, R Amygdala, R ParaHippocamp, R Insula, R Front Med Orb, R Subgenual, R Rolandic Oper, R Front Inf Ope, R Front Mid,* and *R Precentral*.

A similar pattern was observed for clustering coefficient. Above-chance clustering-coefficient loss was found in 55 regions and was again biased toward frontal, cingulate, parietal, and subcortical association territories. These regions were: *L Precentral, L Frontal Sup, L Front Sup Orb, L Front Mid, L Front Mid Orb, L Front Inf Ope, L Front Inf Tri, L Front Inf Orb, L Rolandic Oper, L Supp Motor Ar, L Front Sup Med, L Ventr Med PFC, L Front Med Orb, L Insula, L Cingulum Ant, L Cingulum Mid, L Hippocampus, L Occipital Mid, L Postcentral, L Parietal Sup, L Parietal Inf, L SupraMarginal, L Angular, L Precuneus, L Paracentr Lob, L Caudate, L Putamen, L Pallidum, L Thalamus, L Temporal Mid, L Temporal Inf, R Thalamus, R Pallidum, R Putamen, R Caudate, R Paracentr Lob, R Precuneus, R Angular, R Parietal Sup, R Postcentral, R Occipital Mid, R Occipital Sup, R Cingulum Mid, R Cingulum Ant, R Front Med Orb, R Ventr Med PFC, R Front Sup Med, R Supp Motor Ar, R Front Inf Tri, R Front Inf Ope, R Front Mid Orb, R Front Mid, R Front Sup Orb, R Frontal Sup, and R Precentral*.

Regions showing below-chance clustering-coefficient loss were fewer (14 regions) and again emphasised sensory and medial-temporal structures: *L ParaHippocamp, L Amygdala, L Calcarine, L Lingual, L Fusiform, L Heschl, L Tempr Pol Mid, R Tempr Pol Sup, R Heschl, R Parietal Inf, R Occipital Inf, R ParaHippocamp, R Insula, and R Rolandic Oper*.

### Rich-club analysis

At the level of connection classes, Dataset 1 showed a clear gradient in hotspot dependence across the rich-club architecture (**Figure 3C**). A repeated-measures ANOVA revealed a strong main effect of connection type (F(2,154) = 2956.93, p < 1e-123, ηp² = 0.975). Rich–rich connections showed the greatest proportion of routing through the ageing hotspot (mean = 0.528, s.d. = 0.043), followed by rich–periphery connections (mean = 0.448, s.d. = 0.036), whereas periphery–periphery connections showed the lowest dependence (mean = 0.306, s.d. = 0.027). Pairwise comparisons confirmed that rich-rich connections were more hotspot-dependent than both periphery-periphery connections (t(77) = 61.85, p = 2.11e-67), and rich-periphery connections (t(77) = 26.57, p = 1.58e-40); rich-periphery connections were in turn more hotspot-dependent than periphery-periphery connections (t(77) = 72.47, p = 1.30e-72). The same ordering was replicated in Dataset 2, again with a strong main effect of connection type (F(2,174) = 1468.09, p < 1e-100, ηp² = 0.944). Rich-rich connections showed the highest hotspot dependence (mean = 0.221, s.d. = 0.036), followed by rich-periphery connections (mean = 0.158, s.d. = 0.016), and periphery-periphery connections (mean = 0.076, s.d. = 0.009). Pairwise tests showed the same graded pattern, with rich-rich > periphery-periphery (t(87) = 41.30, p = 6.06e-59), rich-rich > rich-periphery (t(87) = 22.82, p = 1.74e-38), and rich-periphery > periphery-periphery (t(87) = 62.34, p = 5.53e-74). Together, these findings indicate that age-sensitive white matter is disproportionately embedded in the brain’s integrative backbone, with rich-club routes showing the strongest dependence on the hotspot tissue.

## Discussion

In this study, we combined TBSS-defined white-matter hotspots with a simulated-lesion connectome framework to ask what age-sensitive white matter means for large-scale structural communication. Across two independent datasets, we found that age-sensitive hotspots reduced structural network integration, but did so in a way that was consistently less disruptive than a matched random lesion. At the same time, these hotspots were not distributed evenly across the connectome. Their effects were concentrated in high-degree nodes, rich-club routes, and frontal–cingulate–subcortical association systems, whereas more peripheral and sensory systems were comparatively spared. Taken together, these findings suggest that healthy ageing is not well described as a uniform weakening of connectivity. Instead, it follows a selective and constrained pattern in which the brain’s integrative core is disproportionately affected, while global communication remains more preserved than would be expected from an equally sized random lesion (Aerts et al., 2016).

The current work builds on recent studies showing that spatial patterns of white-matter damage can be translated into network-level consequences, but extends this logic from white matter hyperintensities (WMH) and small-vessel disease to microstructural hotspots of healthy ageing. Earlier work demonstrated that WMH-related disconnection is associated with lower cognitive performance – especially executive function – and with preferential loss of interhemispheric, long-range, frontal, and subcortical connectivity (Langen et al., 2018; Petersen et al., 2020). Simulated-lesion studies further showed that WMH masks or WMH frequency maps can be used as tractography constraints to estimate disconnectomes linked to cognition (Li et al., 2023; Taghvaei et al., 2024), while simulations of progressive periventricular white-matter ischemia suggested that spatially patterned white-matter injury selectively disrupts structural communication routes (Li et al., 2021). Here, we extend this framework to empirically defined microstructural ageing hotspots, providing a bridge between normative white-matter ageing and the broader disconnectome literature.

A central implication of the efficiency result is that the topology of age-sensitive white-matter change matters, not just its overall extent. In our data, the empirical lesion always reduced global efficiency less than the matched-mass null (**Figure 2B**). We interpret this as evidence for a constrained form of decline: the ageing pattern is clearly damaging, but it is not maximally disruptive from a network perspective. It suggests that the spatial distribution of healthy-ageing white-matter change leaves more global communication capacity intact than alternative patterns with the same total lesion burden. In that sense, the result fits naturally with broader frameworks that distinguish neural vulnerability and decline from maintenance, reserve, and compensation; however, we refrain from behavioural interpretation in the absence of direct cognitive modelling here (Cabeza et al., 2018).

Another major finding is that ageing-sensitive tissue is disproportionately embedded in the brain’s integrative backbone (**Figure 3**). This was visible at several levels of the analysis: the hub-disruption index was positive, meaning that nodes that are more connected in the network lose proportionally more in ageing; nodewise losses were biased toward frontal, cingulate, and subcortical association structures; and rich–rich connections showed the strongest dependence on hotspot tissue in both datasets. This converges with recent structural-connectome work showing that age effects are stronger in hub than in peripheral connections and that ageing-sensitive subnetworks are especially relevant for late-life cognitive decline (Li et al., 2023a).

The spatial pattern of the nodewise results is equally important. Above-chance loss was concentrated in frontal, cingulate, and thalamo-striatal regions, while below-chance loss was more common in posterior sensory and temporo-limbic territories (**Figure 3B**). This broadly matches long-standing diffusion MRI observations that later-myelinating association tracts are more vulnerable than earlier-maturing sensory pathways, in line with retrogenesis or “last-in–first-out” accounts (Brickman et al., 2012). It also agrees with the broader disconnection literature, which has emphasised that age-related cognitive decline is linked to degradation of the white-matter pathways that support efficient communication among distributed systems (Bennett & Madden, 2014; Madden et al., 2017). The novel step here is that we do not stop at the spatial map itself: we show this association-biased vulnerability in a data-driven way and provide evidence for measurable consequences for connectome topology, shifting the emphasis from where ageing occurs to what network roles those affected pathways support.

These findings also help place structural ageing more concretely within broader theories of cognitive ageing. At the most direct level, they support disconnection accounts by showing that age-sensitive white matter is especially tied to the routes that support network integration (Bennett & Madden, 2014; Madden et al., 2017). Despite strong effects on network integration and backbone vulnerability, the relative sparing of peripheral and sensory systems offers a plausible structural context for compensation-based frameworks. In STAC, compensatory scaffolding is thought to rely on the recruitment of alternative circuits when core systems deteriorate (Park & Reuter-Lorenz, 2009). Our findings are compatible with that logic in showing that not all routes are equally compromised, leaving room for rerouting through relatively preserved pathways. Similarly, PASA describes a shift toward greater anterior engagement in older adults during cognitive tasks (Davis et al., 2008). Although the present data are structural rather than functional, they provide a potential structural connectivity counterpart to that shift: when communication through the integrative backbone is selectively stressed, greater reliance on higher-order control systems becomes easier to understand as a systems-level consequence rather than a purely functional phenomenon. However, our study does not test compensation or functional recruitment directly; it defines structural constraints within which such processes may occur.

Methodologically, the study contributes a useful bridge between TBSS and connectomics. Previous work has shown that it is feasible to move from a spatial pattern of white-matter abnormality to network-level consequences, but this has been done mainly in white-matter hyperintensity and small-vessel-disease research rather than in healthy-ageing microstructural hotspots (Langen et al., 2018; Li et al., 2023b). By contrast, our approach starts from data-driven diffusion hotspots and quantifies how strongly individual structural edges depend on those hotspots. The cross-dataset design was important here because it allowed us to separate hotspot discovery from hotspot-constrained tractography and thus avoiding circularity. The matched-mass null added a second layer of inference by testing whether the empirical topology of lesioning was more or less disruptive than would be expected by chance given the same total lesion burden. Together, these elements offer a straightforward, data-driven framework for relating specific spatial patterns of ageing, and potentially other dMRI-derived correlates, to constraints in whole-brain connectivity patterns (Aerts et al., 2016; Roberts et al., 2017).

Grounded on a solid methodological basis, the present work also opens two clear avenues for further development. First, the cross-sectional design allows us to characterise the topology of healthy-ageing differences, but does not yet capture within-person adaptation over time. Extending this framework to longitudinal data will make it possible to test whether the same constrained patterns anticipate subsequent structural or cognitive trajectories. Second, although the study is theoretically grounded in cognition, the current analyses focus on structural organisation and do not yet explicitly link simulated lesion burden to cognitive performance or functional reorganisation. Integrating these domains represents an important next step toward understanding how structural constraints shape maintenance, compensation, and behavioural variability in later life.

These considerations point directly to the broader potential of the framework. Given converging evidence that ageing is accompanied by changes in large-scale functional organisation, altered balance between integration and segregation, and broader neurophysiological reconfiguration across the lifespan (Sala-Llonch et al., 2015; Deery et al., 2023; Fernández-Rubio et al., 2025; Malvaso et al., 2025), a natural extension of this work is to combine the present approach with cognitive and functional measures. This may be especially informative in light of recent findings showing age-related differences in auditory short-term, long-term, and working memory, as well as neural reorganisation during the recognition of complex temporal sequences, suggesting that age-related changes in behaviour and neurophysiology are tightly linked rather than independent phenomena (Fernández-Rubio et al., 2024; Bonetti et al., 2024; Costa et al., 2025). More broadly, recent theoretical and review work has argued that predictive-processing paradigms may offer a particularly sensitive window into ageing-related changes in brain function and plasticity, and may therefore provide a valuable context in which to test whether individuals whose hotspots place greater stress on the integrative backbone also show corresponding shifts in functional organisation or differences in cognitive performance (Heng et al., 2024). This broader translational direction also aligns with growing calls to connect brain-health markers to systems-level function and behavioural outcomes across ageing, both in healthy populations and in the prevention and treatment of neurodegenerative disease (Beard et al., 2016; Sabayan et al., 2023; Livingston et al., 2024). In this sense, the present framework provides a generalisable tool for translating spatial patterns of white-matter disruption into connectome-level consequences across both healthy and pathological ageing.

Taken together, the present findings suggest a reframing of healthy ageing: rather than reflecting uniform connectivity loss, it appears as a process of selective and constrained reconfiguration, in which age-sensitive white matter disproportionately impacts the brain’s integrative core while preserving more communication capacity than would be expected under random decline.

## Materials and methods

The goal of the study was to move from where age-related white-matter differences are observed to what those differences mean for large-scale structural communication. To do so, we combined voxel-wise hotspot detection with connectome analysis in two independent datasets. We identify age-sensitive white-matter clusters in one dataset and test their network consequences in the other, thereby reducing circularity when estimating how strongly the observed hotspots constrain whole-brain connectivity. We then benchmark those effects against a matched-mass null to increase interpretability.

### Participants and study design

We analysed diffusion MRI (dMRI) data from two independent datasets of healthy young and older adults (Dataset 1: n = 77; 37 young adults, mean age = 21.89 years, s.d. = 2.05; 40 older adults, mean age = 67.50 years, s.d. = 5.46; ethical approval by the Institutional Review Board at Aarhus University [Case No. DNC-IRB-2021-012]; Dataset 2: n = 67; 41 young adults, mean age = 22.37 years, s.d. = 2.57; 26 older adults, mean age = 68.77 years, s.d. = 5.30; ethical approval by the Ethics Committee of the Central Denmark Region [De Videnskabsetiske Komitéer for Region Midtjylland, Ref 1-10-72-127-23]). The experimental procedures were carried out in compliance with the Declaration of Helsinki – Ethical Principles for Medical Research. All participants gave informed consent before starting the experimental procedure. Both datasets were processed with the same preprocessing, TBSS, and tractography pipeline. The key design feature was a cross-dataset hotspot framework: age-sensitive white-matter clusters were identified separately within each dataset, but were used as tractography waypoints in the other dataset to avoid circularity when estimating network consequences.

### Diffusion preprocessing and TBSS

For both datasets, diffusion data were converted from DICOM to NIfTI together with gradient tables. Susceptibility-induced distortions were corrected with TOPUP using paired AP/PA b0 images, brain masks were derived from the corrected mean b0 image, and eddy-current distortions and head motion were corrected with EDDY while incorporating the TOPUP field estimates. Diffusion tensors were then fitted voxelwise to obtain standard scalar maps, and fractional anisotropy (FA) served as the primary microstructural measure (Basser et al., 1994; Andersson & Sotiropoulos, 2016).

TBSS was performed separately in each dataset (Smith et al., 2006). FA maps were nonlinearly registered to standard space, projected onto the mean FA skeleton, and thresholded at FA = 0.2. Older and younger adults were compared voxelwise with two-sample tests. Significant age-sensitive clusters were defined with a permutation-based cluster-extent procedure (10,000 label shuffles; cluster-forming threshold p < .005, two-sided), and the union of suprathreshold voxels was saved as a binary hotspot mask in MNI space for each dataset (**Figure 1A**).

### Probabilistic tractography and connectome construction

For each participant, the AAL-90 atlas and the relevant hotspot mask were transformed into diffusion space using subject-specific registrations, with nearest-neighbour interpolation for label images (Tzourio-Mazoyer et al., 2002). Local fibre orientations were estimated with BEDPOSTX, and probabilistic tractography was run with PROBTRACKX2 in network mode using 5,000 streamlines per seed (Behrens et al., 2003, 2007, Bruzzone et al., 2022) (**Figure 1A**).

Two connectivity matrices were reconstructed per participant. The first was a raw structural connectome obtained without any waypoint constraint (Mraw). The second was a cluster-through connectome in which only streamlines passing through the age-sensitive hotspot were counted (Mclust) (**Figure 1B**). To avoid circularity, cluster-through tractography was always performed cross-dataset: hotspots identified in Dataset 1 were used in Dataset 2, and vice versa (**Figure 1D**). Edge weights were symmetrised and normalised by the sum of the two endpoint ROI volumes in diffusion space to correct for head-size and ROI-size biases.

Because probabilistic tractography yields dense matrices with many weak edges, networks were pruned using consistency-based thresholding (Roberts et al., 2017). Edge consistency was defined across participants from the raw matrices as the ratio of standard deviation to mean, and 30% of the most consistent edges were retained. The same edge mask was applied to the corresponding raw and cluster-through matrices within each dataset.

### Simulated lesion

For each participant, hotspot dependence was quantified by comparing the raw and cluster-through matrices. This yielded an edgewise, cluster-proportional simulated lesion in which each connection was down-weighted according to how much of its reconstructed streamline weight traversed the age-sensitive tissue. The resulting lesioned network therefore reflected empirical hotspot routing rather than a binary or abstract attack (**Figure 1B**).

### Permutation framework

Observed effects were benchmarked against a matched-mass null model. For each participant, we generated up to 10,000 random lesions that preserved the total lesion mass of the empirical lesion while redistributing that removal across eligible edges. This yielded participant-specific null distributions for each graph metric under a random lesion with the same overall weight.

### Graph measures

We quantified weighted global efficiency, nodewise clustering coefficient, and weighted degree for the raw and lesioned networks using standard graph-theoretical measures (Latora & Marchiori, 2001; Rubinov & Sporns, 2010; Onnela et al., 2005) (**Figure 1C**). For each participant, empirical lesion effects were defined as the pre- to post-lesion change in these measures.

### Global efficiency

For each participant, the observed decrease in global efficiency was compared with the matched-mass permutation distribution to obtain a one-sided p-value. This tested whether the empirical hotspot-constrained lesion reduced network integration more than expected from a random lesion of equal size.

### Nodewise degree and clustering coefficient

For each participant and node, the observed decreases in degree and clustering coefficient were compared with the corresponding matched-mass permutation distributions, yielding one-sided p-values for greater-than-chance loss and for relative sparing. To identify effects that were consistent across the full sample, participant-level p-values from both datasets were pooled separately for degree and clustering coefficient. For each node, the balance of above-chance loss versus below-chance loss across participants was then tested with a binomial model, followed by false-discovery-rate correction across the 90 regions. This produced group-level maps of nodes showing reliable above-chance loss or reliable relative sparing.

### Hub-disruption analysis

To test whether highly connected nodes were disproportionately affected, proportional degree loss was regressed against pre-lesion degree within each participant. The resulting slope was used as a hub-disruption index, with more positive values indicating stronger proportional loss in high-degree nodes.

### Rich-club analysis

Rich-club organisation was assessed using weighted rich-club coefficients relative to degree-preserving random graphs (van den Heuvel & Sporns, 2011). Rich-club nodes were identified at degree thresholds where the normalised coefficient exceeded 1, and edges were classified as rich-rich, rich-periphery, or periphery-periphery. For each edge class, hotspot dependence was quantified as the proportion of streamline weight routed through age-sensitive tissue, and these values were compared statistically across classes.

## Data availability

The neuroimaging data related to the experiment is available upon reasonable request.

## Code availability

The codes are available at the following link: https://github.com/cognikita-cn/DTI_HealthyAgeing.git

## Acknowledgements

The Center for Music in the Brain (MIB) is funded by the Danish National Research Foundation (project number DNRF117), The Lundbeck Foundation (R469-2024-1573) and Købmand Herman Sallings Fond.

L.B. is supported by Sapere Aude: Independent Research Fund Denmark (DFF) Research Leader (grant ID: 10.46540/5253-00003B), Lundbeck Foundation (Talent Prize 2022), Carlsberg Foundation (CF20-0239), Center for Music in the Brain, Linacre College of the University of Oxford and Nordic Mensa Fund.

MLK is supported by Center for Music in the Brain and Centre for Eudaimonia and Human Flourishing, which is funded by the Pettit and Carlsberg Foundations.

We thank Emma Risgaard Olsen, Mathias Klarlund, Orla Mallon, Francesco Carlomagno, Mathias H. Andersen, Luna Frausing and Victor Pando-Naude for their important contribution during the data collection, and Giancarlo Valente for his mentorship during this work.

## Author contributions

N.K. conceived the initial hypotheses and methodological development, which were further refined by L.B. and M.R. Resources were recruited by L.B., M.L.K., E.S. and P.V. for performing data collection and analysis. G.F.R., E.S. and L.B. collected the data. N.K. and L.B. performed pre-processing. N.K. developed the simulated lesion approach and statistical analysis. M.L.K., P.V., L.B. and M.R. provided essential help to interpret and frame the results within the neuroscientific and analytical literature. N.K. wrote the first draft of the manuscript, which was primarily integrated by L.B. and M.R. The figures were prepared by N.K., with the support of L.B. and M.R. All the authors contributed to and approved the final version of the manuscript.

## References

Aerts, H., Fias, W., Caeyenberghs, K., & Marinazzo, D. (2016). Brain networks under attack: Robustness properties and the impact of lesions. Brain, 139(12), 3063–3083. 10.1093/brain/aww194

Andersson, J. L. R., & Sotiropoulos, S. N. (2016). An integrated approach to correction for off-resonance effects and subject movement in diffusion MR imaging. NeuroImage, 125, 1063–1078. 10.1016/j.neuroimage.2015.10.019

Barrick, T. R., Charlton, R. A., Clark, C. A., & Markus, H. S. (2010). White matter structural decline in normal ageing: A prospective longitudinal study using tract-based spatial statistics. NeuroImage, 51(2), 565–577. 10.1016/j.neuroimage.2010.02.033

Basser, P. J., Mattiello, J., & LeBihan, D. (1994). MR diffusion tensor spectroscopy and imaging. Biophysical Journal, 66(1), 259–267. 10.1016/S0006-3495(94)80775-1

Beard, J. R., Officer, A., de Carvalho, I. A., Sadana, R., Pot, A. M., Michel, J.-P., Lloyd-Sherlock, P., Epping-Jordan, J. E., Peeters, G. M. E. E., Mahanani, W. R., Thiyagarajan, J. A., & Chatterji, S. (2016). The World report on ageing and health: A policy framework for healthy ageing. The Lancet, 387(10033), 2145–2154. 10.1016/S0140-6736(15)00516-4

Behrens, T. E. J., Woolrich, M. W., Jenkinson, M., Johansen-Berg, H., Nunes, R. G., Clare, S., Matthews, P. M., Brady, J. M., & Smith, S. M. (2003). Characterization and propagation of uncertainty in diffusion-weighted MR imaging. Magnetic Resonance in Medicine, 50(5), 1077–1088. 10.1002/mrm.10609

Behrens, T. E., Berg, H. J., Jbabdi, S., Rushworth, M. F., & Woolrich, M. W. (2007). Probabilistic diffusion tractography with multiple fibre orientations: What can we gain?. NeuroImage, 34(1), 144–155. 10.1016/j.neuroimage.2006.09.018

Bennett, I. J., & Madden, D. J. (2014). Disconnected aging: Cerebral white matter integrity and age-related differences in cognition. Neuroscience, 276, 187–205. 10.1016/j.neuroscience.2013.11.026

Bonetti, L., Fernández-Rubio, G., Lumaca, M., Carlomagno, F., Risgaard Olsen, E., Criscuolo, A., Kotz, S. A., Vuust, P., Brattico, E., & Kringelbach, M. L. (2024). Age-related neural changes underlying long-term recognition of musical sequences. Communications Biology, 7(1), Article 1036. 10.1038/s42003-024-06587-7

Bruzzone, S.E.P., Lumaca, M., Brattico, E. et al. Dissociated brain functional connectivity of fast versus slow frequencies underlying individual differences in fluid intelligence: a DTI and MEG study. Sci Rep 12, 4746 (2022). 10.1038/s41598-022-08521-5

Brickman, A. M., Meier, I. B., Korgaonkar, M. S., Provenzano, F. A., Grieve, S. M., Siedlecki, K. L., Wasserman, B. T., Williams, L. M., & Zimmerman, M. E. (2012). Testing the white matter retrogenesis hypothesis of cognitive aging. Neurobiology of Aging, 33(8), 1699–1715. 10.1016/j.neurobiolaging.2011.06.001

Cabeza, R., Albert, M., Belleville, S., Craik, F. I. M., Duarte, A., Grady, C. L., Lindenberger, U., Nyberg, L., Park, D. C., Reuter-Lorenz, P. A., Rugg, M. D., Steffener, J., & Rajah, M. N. (2018). Maintenance, reserve and compensation: The cognitive neuroscience of healthy ageing. Nature Reviews Neuroscience, 19(11), 701–710. 10.1038/s41583-018-0068-2

Coelho, A., Fernandes, H. M., Magalhães, R., Moreira, P. S., Marques, P., Soares, J. M., … & Sousa, N. (2021). Signatures of white-matter microstructure degradation during aging and its association with cognitive status. Scientific reports, 11(1), 4517.

Costa, M., P. Vuust, M. L. Kringelbach, and L. Bonetti. 2025. “EEG Correlates of Auditory Short-Term Memory and Dissimilarity Perception in Young and Older Adults.” European Journal of Neuroscience61, no. 12: e70166. 10.1111/ejn.70166.

Davis, S. W., Dennis, N. A., Daselaar, S. M., Fleck, M. S., & Cabeza, R. (2008). Que PASA? The posterior–anterior shift in aging. Cerebral Cortex, 18(5), 1201–1209. 10.1093/cercor/bhm155

Deery, H. A., Di Paolo, R., Moran, C., Egan, G. F., & Jamadar, S. D. (2023). The older adult brain is less modular, more integrated, and less efficient at rest: A systematic review of large-scale resting-state functional brain networks in aging. Psychophysiology, 60(1), e14159. 10.1111/psyp.14159

Fernández-Rubio, G., Olsen, E. R., Klarlund, M., Mallon, O. P., Carlomagno, F., Vuust, P., Kringelbach, M. L., Brattico, E., & Bonetti, L. (2024). Investigating the impact of age on auditory short-term, long-term, and working memory. Psychology of Music, 52(2), 187–198. 10.1177/03057356231183404

Fernández-Rubio, G., Vuust, P., Kringelbach, M. L., & Bonetti, L. (2025). The neurophysiology of healthy and pathological aging: A comprehensive systematic review. Brain Structure and Function, 230(8), Article 146. 10.1007/s00429-025-03012-5

GBD 2019 Dementia Forecasting Collaborators. (2022). Estimation of the global prevalence of dementia in 2019 and forecasted prevalence in 2050: An analysis for the Global Burden of Disease Study 2019. The Lancet Public Health, 7(2), e105–e125. 10.1016/S2468-2667(21)00249-8

Gunning-Dixon, F. M., & Raz, N. (2000). The cognitive correlates of white matter abnormalities in normal aging: A quantitative review. Neuropsychology, 14(2), 224– 232. 10.1037/0894-4105.14.2.224

Gunning-Dixon, F. M., Brickman, A. M., Cheng, J. C., & Alexopoulos, G. S. (2009). Aging of cerebral white matter: A review of MRI findings. International Journal of Geriatric Psychiatry, 24(2), 109–117. 10.1002/gps.2087

Heng, J. G., Zhang, J., Bonetti, L., Lim, W. P. H., Vuust, P., Agres, K., & Chen, S. H. A. (2024). Understanding music and aging through the lens of Bayesian inference. Neuroscience and Biobehavioral Reviews, 163, Article 105768. 10.1016/j.neubiorev.2024.105768

Jones, D. K., Knösche, T. R., & Turner, R. (2013). White matter integrity, fiber count, and other fallacies: The do’s and don’ts of diffusion MRI. NeuroImage, 73, 239–254. 10.1016/j.neuroimage.2012.06.081

Langen, C. D., Cremers, L. G. M., de Groot, M., White, T., Ikram, M. A., Niessen, W. J., & Vernooij, M. W. (2018). Disconnection due to white matter hyperintensities is associated with lower cognitive scores. NeuroImage, 183, 745–756. 10.1016/j.neuroimage.2018.08.037

Latora, V., & Marchiori, M. (2001). Efficient behavior of small-world networks. Physical Review Letters, 87(19), 198701. 10.1103/PhysRevLett.87.198701

Li, M., Habes, M., Grabe, H., Kang, Y., Qi, S., & Detre, J. A. (2023). Disconnectome associated with progressive white matter hyperintensities in aging: A virtual lesion study. Frontiers in Aging Neuroscience, 15, Article 1237198. 10.3389/fnagi.2023.1237198

Li, X., Salami, A., & Persson, J. (2023). Hub architecture of the human structural connectome: Links to aging and processing speed. NeuroImage, 278, Article 120270. 10.1016/j.neuroimage.2023.120270

Li, Z., Dolui, S., Habes, M., Bassett, D. S., Wolk, D., & Detre, J. A. (2021). Predicted disconnectome associated with progressive periventricular white matter ischemia. Cerebral Circulation - Cognition and Behavior, 2, Article 100022. 10.1016/j.cccb.2021.100022

Livingston, G., Huntley, J., Liu, K. Y., Costafreda, S. G., Selbæk, G., Alladi, S., Ames, D., Ballard, C., Banerjee, S., Burns, A., Cohen-Mansfield, J., Dickson, C., Farina, N., Ferri, C. P., Gitlin, L. N., Howard, R., Kales, H. C., Kivimäki, M., Larson, E. B., … Mukadam, N. (2024). Dementia prevention, intervention, and care: 2024 report of the Lancet Standing Commission. The Lancet, 404(10452), 572–628. 10.1016/S0140-6736(24)01296-0

Madden, D. J., Bennett, I. J., Burzynska, A., Potter, G. G., Chen, N.-K., & Song, A. W. (2017). Sources of disconnection in neurocognitive aging: Cerebral white-matter integrity, resting-state functional connectivity, and white-matter hyperintensity volume. Neurobiology of Aging, 54, 199–213. 10.1016/j.neurobiolaging.2017.01.027

Madole, J. W., Ritchie, S. J., Cox, S. R., Buchanan, C. R., Valdés Hernández, M. C., Muñoz Maniega, S., Wardlaw, J. M., Harris, M. A., Bastin, M. E., Deary, I. J., & Tucker-Drob, E. M. (2021). Aging-sensitive networks within the human structural connectome are implicated in late-life cognitive declines. Biological Psychiatry, 89(8), 795–806. 10.1016/j.biopsych.2020.06.010

Malvaso, C., Fernández-Rubio, G., Rosso, M., Serra, E., Rudi, V., Vuust, P., … & Bonetti, L. (2025). FREQ-NESS reveals age-related differences in frequency-resolved brain networks during auditory recognition and resting state. bioRxiv, 2025–06.

Onnela, J.-P., Saramäki, J., Kertész, J., & Kaski, K. (2005). Intensity and coherence of motifs in weighted complex networks. Physical Review E, 71(6), 065103. 10.1103/PhysRevE.71.065103

Park, D. C., & Reuter-Lorenz, P. A. (2009). The adaptive brain: Aging and neurocognitive scaffolding. Annual Review of Psychology, 60, 173–196. 10.1146/annurev.psych.59.103006.093656

Peter, C., Sathe, A., Shashikumar, N., Pechman, K. R., Workmeister, A. W., Jackson, T. B., Huo, Y., Mukherjee, S., Mez, J., Dumitrescu, L., Gifford, K. A., Bolton, C. J., Gaynor, L. S., Risacher, S. L., Beason-Held, L. L., An, Y., Arfanakis, K., Erus, G., Davatzikos, C., … Tosun-Turgut, D. (2025). White matter abnormalities and cognition in aging and Alzheimer disease. JAMA Neurology, 82(8), 825–836. 10.1001/jamaneurol.2025.1601

Petersen, M., Frey, B. M., Schlemm, E., Mayer, C., Hanning, U., Thomalla, G., … Fiehler, J. (2020). Network localisation of white matter damage in cerebral small vessel disease. Scientific Reports, 10, Article 9210. 10.1038/s41598-020-66013-w

Poulakis, K., Reid, R. I., Przybelski, S. A., Knopman, D. S., Graff-Radford, J., Lowe, V. J., Mielke, M. M., Machulda, M. M., Jack, C. R., Jr., Petersen, R. C., Westman, E., & Vemuri, P. (2021). Longitudinal deterioration of white-matter integrity: Heterogeneity in the ageing population. Brain Communications, 3(1), fcaa238. 10.1093/braincomms/fcaa238

Prince, J. B., Davis, H. L., Tan, J., Muller-Townsend, K., Markovic, S., Lewis, D. M. G., Hastie, B., Thompson, M. B., Drummond, P. D., Fujiyama, H., & Sohrabi, H. R. (2024). Cognitive and neuroscientific perspectives of healthy ageing. Neuroscience and biobehavioral reviews, 161, 105649. 10.1016/j.neubiorev.2024.105649

Riedel, L., van den Heuvel, M. P., & Markett, S. (2022). Trajectory of rich club properties in structural brain networks. Human Brain Mapping, 43(14), 4239–4253. 10.1002/hbm.25950

Roberts, J. A., Perry, A., Roberts, G., Mitchell, P. B., & Breakspear, M. (2017). Consistency-based thresholding of the human connectome. NeuroImage, 145, 118– 129. 10.1016/j.neuroimage.2016.09.053

Rubinov, M., & Sporns, O. (2010). Complex network measures of brain connectivity: Uses and interpretations. NeuroImage, 52(3), 1059–1069. 10.1016/j.neuroimage.2009.10.003

Sabayan, B., Doyle, S., Rost, N. S., Sorond, F. A., Lakshminarayan, K., & Launer, L. J. (2023). The role of population-level preventive care for brain health in ageing. The Lancet Healthy Longevity, 4(6), e274–e283. 10.1016/S2666-7568(23)00051-X

Sala-Llonch, R., Bartrés-Faz, D., & Junqué, C. (2015). Reorganization of brain networks in aging: A review of functional connectivity studies. Frontiers in Psychology, 6, Article 663. 10.3389/fpsyg.2015.00663

Sexton, C. E., Walhovd, K. B., Storsve, A. B., Tamnes, C. K., Westlye, L. T., Johansen-Berg, H., & Fjell, A. M. (2014). Accelerated changes in white matter microstructure during aging: a longitudinal diffusion tensor imaging study. The Journal of neuroscience: the official journal of the Society for Neuroscience, 34(46), 15425–15436. 10.1523/JNEUROSCI.0203-14.2014

Smith, S. M., Jenkinson, M., Johansen-Berg, H., Rueckert, D., Nichols, T. E., Mackay, C. E., Watkins, K. E., Ciccarelli, O., Cader, M. Z., Matthews, P. M., & Behrens, T. E. J. (2006). Tract-based spatial statistics: Voxelwise analysis of multi-subject diffusion data. NeuroImage, 31(4), 1487–1505. 10.1016/j.neuroimage.2006.02.024

Stumme, J., Krämer, C., Miller, T., Schreiber, J., Caspers, S., & Jockwitz, C. (2022). Interrelating differences in structural and functional connectivity in the older adult’s brain. Human Brain Mapping, 43(18), 5543–5561. 10.1002/hbm.26030

Taghvaei, M., Mechanic-Hamilton, D. J., Sadaghiani, S., Shakibajahromi, B., Dolui, S., Das, S., Brown, C., Gaussoin, S. A., Wang, D. J. J., Detre, J. A., & Habes, M. (2024). Impact of white matter hyperintensities on structural connectivity and cognition in cognitively intact ADNI participants. Neurobiology of Aging, 135, 79– 90. 10.1016/j.neurobiolaging.2023.10.012

Tzourio-Mazoyer, N., Landeau, B., Papathanassiou, D., Crivello, F., Etard, O., Delcroix, N., Mazoyer, B., & Joliot, M. (2002). Automated anatomical labeling of activations in SPM using a macroscopic anatomical parcellation of the MNI MRI single-subject brain. NeuroImage, 15(1), 273–289. 10.1006/nimg.2001.0978

United Nations, Department of Economic and Social Affairs, Population Division. (2024). World population ageing 2023. United Nations.

van den Heuvel, M. P., & Sporns, O. (2011). Rich-club organization of the human connectome. The Journal of Neuroscience, 31(44), 15775–15786. 10.1523/JNEUROSCI.3539-11.2011

Zhao, T., Cao, M., Niu, H., Zuo, X.-N., Evans, A., He, Y., Dong, Q., & Shu, N. (2015). Age-related changes in the topological organization of the white matter structural connectome across the human lifespan. Human Brain Mapping, 36(10), 3777–3792. 10.1002/hbm.22877

